# Sensitive detection of Live *Escherichia coli* by bacteriophage amplification-coupled immunoassay on the Luminex^®^ MAGPIX instrument

**DOI:** 10.1101/318071

**Authors:** Tomotaka Mido, Eric M. Schaffer, Robert W. Dorsey, Shanmuga Sozhamannan, E. Randal Hofmann

## Abstract

Phages are natural predators of bacteria and have been exploited in bacterial detection because of their exquisite specificity to their cognate bacterial hosts. In this study, we present a bacteriophage amplification-coupled assay as a surrogate for detecting a bacterium present in a sample. The assay entails detection of progeny phage resulting from infection and subsequent growth inside the bacterium present in suspected samples. This approach reduces testing time and enhances sensitivity to identify pathogens compared to traditional overnight plaque assay. Further, the assay has the ability to discriminate between live and dead cells since phages require live host cells to infect and replicate. To demonstrate its utility, phage MS2 amplification-coupled, bead-based sandwich type immunoassay on the Luminex^®^ MAGPIX instrument for *Escherichia coli* detection was performed. The assay not only showed live cell discrimination ability but also a limit of *E. coli* detection of 1×10^2^ cells/mL of live cells after a 3-hour incubation. In addition, the sensitivity of the assay was not impaired in the presence of dead cells. These results demonstrate that bacteriophage amplification-coupled assay can be a rapid live cell detection assay compared to traditional culture methods and a promising tool for quick validation of bacterial inactivation. Combined with the unique multiplex bead chemistry afforded by Luminex^®^ MAGPIX platform, the phage assay can be expanded to be an ultra-deep multiplex assay for the simultaneous detection of multiple pathogens using specific phages directed against the target pathogens.

## Introduction

Early diagnosis of an etiological agent is paramount in implementing timely and appropriate countermeasures to prevent fatal consequences. In an outbreak scenario, protecting the patients and preventing further dissemination of the disease relies on early, rapid, accurate and sensitive detection of the infectious agent. This in turn relies on the assay and detection platform used.

Currently, four broad categories of biodetection systems are available. 1) Microbiological/ biochemical tests, 2) antibody based, 3) nucleic acid based and 4) other methods including mass spec and bioluminescence. The length of assay times and levels of purification of the sample to be tested vary widely with these systems. Conventional microbiological culturing and staining, differential growth of target organisms in selective media require live cells and take time anywhere from 16 hours to several days in some cases, prior to definitive identification of the culprit organisms (1).

There are some drawbacks with the antibody or nucleic acid based systems. For example, PCR and nucleic acid sequence-based amplification (NASBA) enrich a single specific piece of DNA or RNA sequence up to 10^6^-fold in 20 minutes to a few hours and theoretically have a sensitivity of a single bacterial cell. The PCR methods give rapid, specific detection but are limited by small sample volumes (e.g., 5 μl for PCR). Furthermore, substances in the sample matrix may inhibit the PCR reaction and the steps used to concentrate the sample to obtain enough templates for PCR may concentrate the inhibitors as well. Immunoassays are based on the concept that any compound that is capable of triggering an immune response can be targeted as an antigen and have been used not only for all types of agents including spores, toxins, and viruses. In general, PCR is much more sensitive than immuno-assays (1). These NASBA and immunoassays cannot discriminate between live and dead target pathogens.

There is another paradigm that takes advantage of phages for bacterial detection. Phages are bacterial viruses and are specific to each bacterial species they infect and sometimes, even strains of a given species. The kinetics of interaction between bacteria and their cognate phages is comparable to that of antigen-antibody interaction, making them highly suitable for bacterial detection (2, 3). In addition, the phage-bacterial specificity has evolved over millions of years making them as good as or even better than antigen-antibody specificity. The specificity is attributed to a receptor on the surface of phage that interacts with a receptor on the bacterial surface and this pair is unique. This specificity has been used to develop phage-typing schemes for bacterial species and strains (4–9). Moreover, the cost incurred in producing a phage-based detection reagent is relatively inexpensive compared to the antigen-antibody based reagents. In addition, phages can be useful in deciphering viability of a bacterial pathogen in the sample and furthermore, replication of phage inside the bacterium leads to an amplification of the detection signals thus increasing the sensitivity of the assay.

A number of detection systems exploiting phage-bacterial specificity have been developed for different bacteria (10). One of the earliest phage based detection systems involved incorporation of *lux* genes in a mycobacterial phage genome. Expression of the *lux* genes in susceptible mycobacterial cells emitted luminescence signals captured by a handheld Polaroid camera device termed “bronx-box” (11). Similar approaches have been taken for construction of recombinant phages for the detection of *Bacillus anthracis* and *Yersinia pestis* (12, 13). Another elegant fluorescence technique, designed to detect deadly *E. coli O157:H7* bacteria, relied on introducing green fluorescent protein (*gfp*) gene via a bacteriophage. Expression of phage-encoded *gfp* inside the bacterium emits fluorescence that can be measured in a flow cytometer (14). These methods involved extensive genetic manipulation and relatively expensive fluorescent measurement instruments. There are other limitations to this approach: a) level of expression of LUX/GFP is dependent on the phage promoter that controls its expression; b) low photostability of GFP permits fluorescence measurement only for a few seconds to a minute under normal microscopic conditions and therefore, renders the quantitative fluorescence assay difficult in GFP expressed cells. In order to improve the sensitivity and potential for multiplexing, phage-quantum dot assays for rapid high-sensitive detection of bacterial pathogens have been described (15, 16). Although phage-quantum dot approach has certain advantages in multiplexing and increased sensitivity, appropriate instruments for measuring multiplex fluorescence signals are not available and thus are not field deployable.

Previous studies have demonstrated the utility of phage amplification coupled-detection assay in a simple platform such as lateral flow immunoassay (LFI) and showed reasonable detection limits (17, 18). Recently, a magnetic bead coupled to phage tail fiber protein has been used as a sensitive tool for detection of *Salmonella* cells (19). Here, we have harnessed phage features with the multiplex capability of MAGPIX platform to develop a phage amplification coupled assay to detect viable bacteria. These features are: 1) The exclusivity of phage infection of live cells; 2) Phage growth following infection resulting in an exponential increase of progeny particles by several orders of magnitude thereby increasing the sensitivity of the assay; 3) The relatively rapid nature of the assay compared to traditional plaque assay or even conventional culture methods. 4) An unparalleled multiplex capability offered by MAGPIX platform because of its unique bead chemistry (20). As a proof of concept, we describe a MAGPIX bead based sandwich type immunoassay, hereafter referred as phage MAGPIX assay, using phage MS2 and anti MS2 antibodies as a surrogate for the detection of *E. coli*. Infection, subsequent replication and growth of MS2 inside *E. coli* present in a sample results in the release of increased number of (several orders of magnitude) progeny MS2 particles, which are captured by the anti MS2 antibody coupled to MAGPIX beads. A secondary (detector) anti-MS2 antibody is added to the complex followed by an additional incubation with streptavidin-coated phycoerythrin (SAPE). The resulting fluorescence of the complex is measured and reported as an indicator of the specific bacteria present in the sample.

## Results

### Determination of assay linearity of phage MS2 based MAGPIX immunoassay

The capture sandwich immunoassay for MS2 phage on the MAGPIX platform was developed using polyclonal anti MS2 antibodies. In the assay, target antigen (MS2) is captured by antibodies (anti MS2 antibodies) coupled on the surface of beads, followed by quantification of the bead bound complex by labeled antibodies. Thus, an increase in progeny phage; i.e., phage amplification can be correlated with amplification in fluorescence signal. The concept of using MAGPIX instrument for this assay is illustrated in Figure 1.

**Figure 1.**
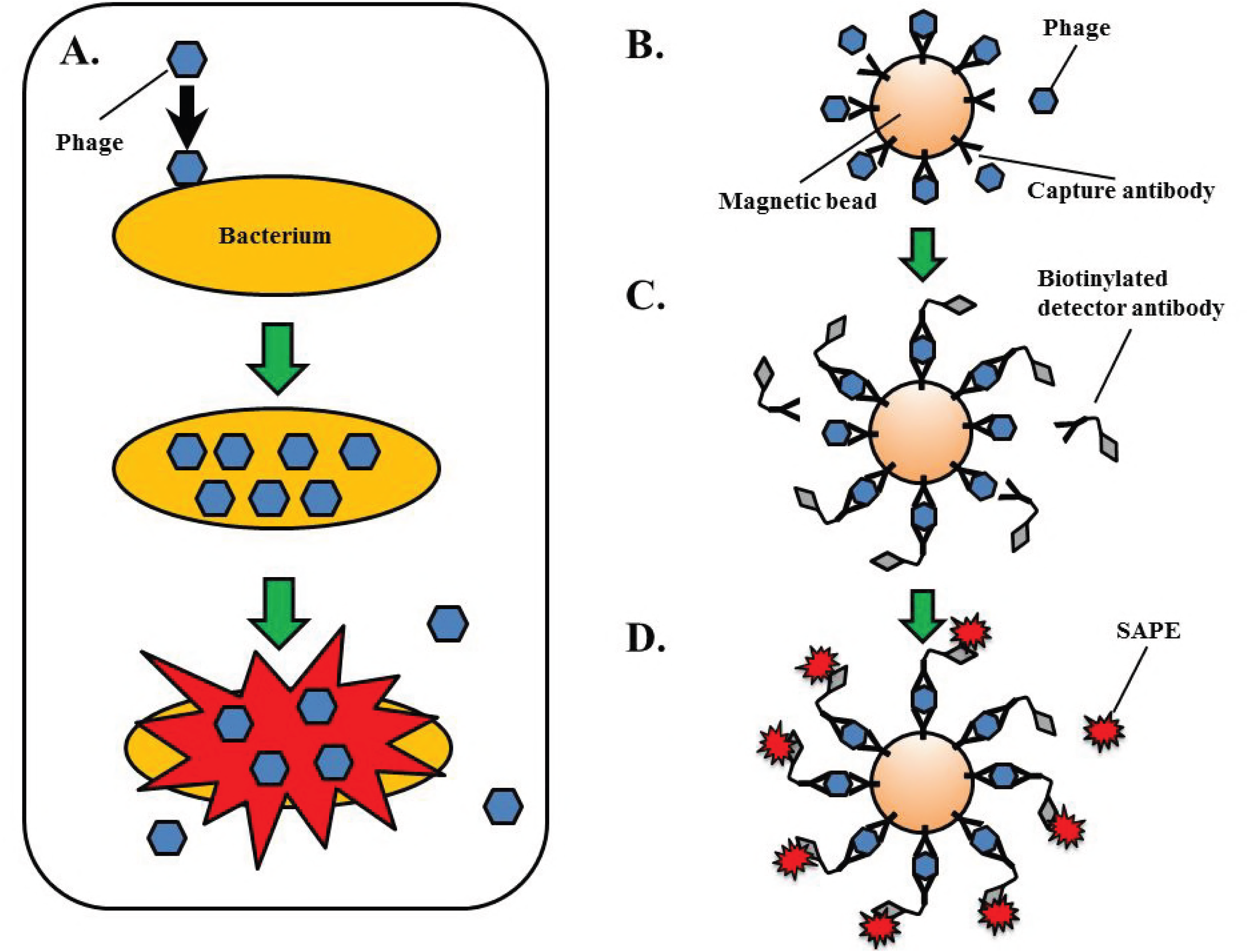
Illustration of bacteriophage amplification-coupled immunoassay for bacterial detection on MAGPIX instrument. (A) Phage infection of a bacterial cell and subsequent growth inside the bacterium results in host cell lysis and release of progeny phage in large numbers. (B) Antibody coupled on the surface of magnetic bead captures input and progeny phage. (C) Addition of detector antibody followed by (D) an incubation with SAPE. The fluorescence emitted by the final complex is measured in the MAGPIX instrument and reported as relative fluorescence units.

The phage MAGPIX assay, as an indirect measurement of bacterial detection, is incumbent upon detection of progeny phages rather than the input phage used to initiate infection. Therefore, the input phage concentration should be low enough (below the detection limits of the instrument) so that upon phage amplification there is high enough phage titer to result in significant signal amplification that can be detected. Also, it should not be too low, in which case the assay would require longer incubation times to produce high enough phage titers that would generate measureable fluorescence signal intensities. In order to determine the appropriate initial MS2 concentration, the linearity of the MS2 MAGPIX assay was assessed. A dose response curve of the assay was generated by serial dilution of MS2 in LB media and measuring the median fluorescence intensity (MFI). A clear linearity of signal intensity was seen at phage concentrations ranging from 1×10^6^ pfu/ mL to 1×10^9^ pfu/ mL (Figure 2). Thus, initial MS2 concentration to assess the signal amplification in the assay based on phage replication was determined to be 1×10^6^ pfu/mL.

**Figure 2.**
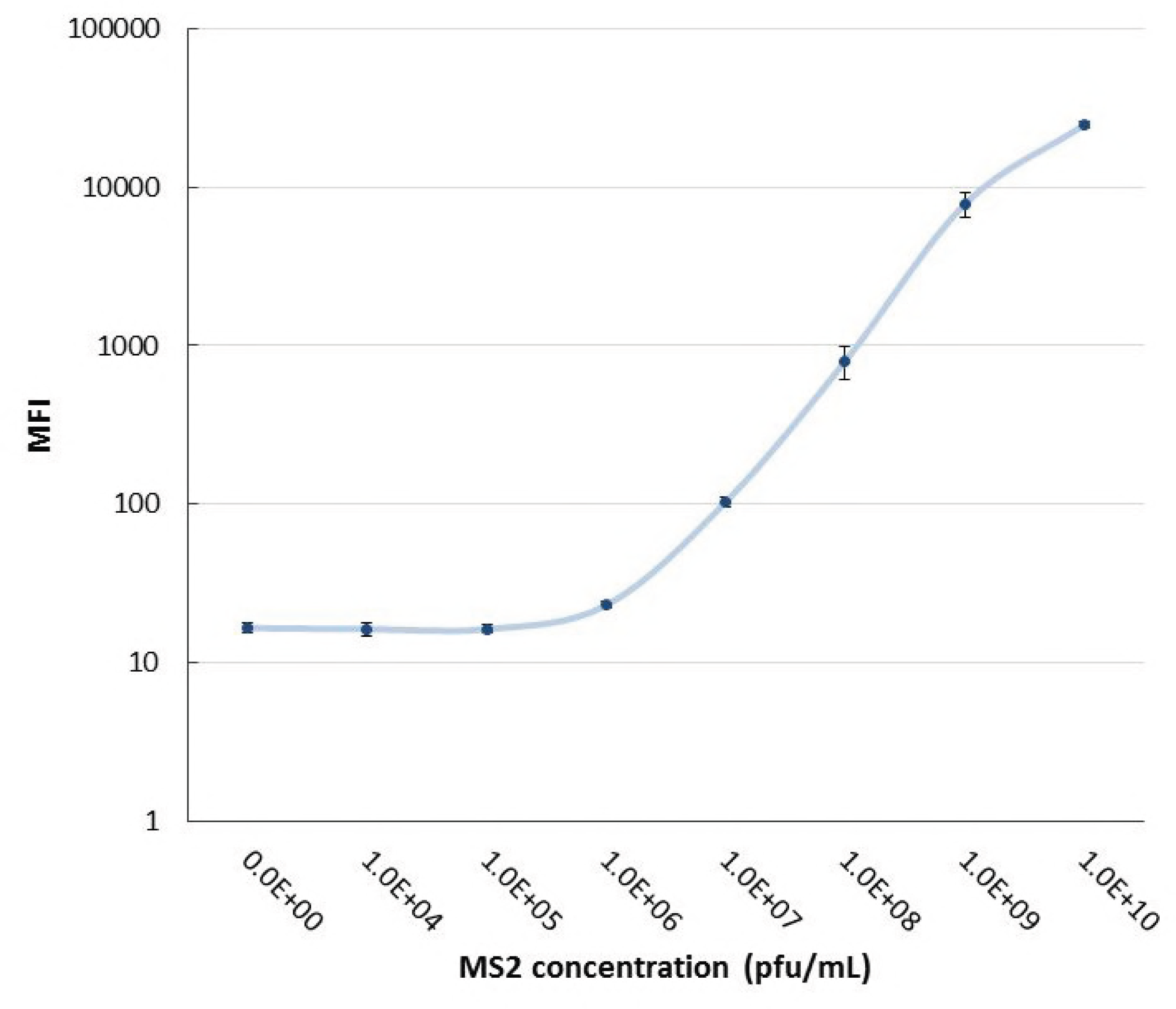
Assay linearity of MS2 MAGPIX immunoassay detection. Sandwich type MAGPIX immunoassay for MS2 detection was performed at respective MS2 concentrations. Vertical axis shows the median fluorescence intensity (MFI). Values are average of two independednt measurements of two replicates each.

### Determination of the live cell discrimination ability of phage MAGPIX immunoassay

In order to evaluate the utility of MS2 amplification-coupled assay for *E. coli* detection, initially, live cell discrimination ability of the assay was assessed. Live cells or heat inactivated *E. coli* cells at a concentration of 1×10^6^ cells/mL were infected with MS2 at a multiplicity of infection of 1 (1×10^6^ pfu/mL) and incubated for 18 hours and the resulting phage particles were analyzed by MS2 MAGPIX immunoassay. The results showed that live cells infected with MS2 amplified fluorescence signal intensity almost 1000 fold at the end of the incubation period whereas dead cells or live cells without the addition of MS2 did not (Figure 3) indicating the ability of the assay to detect live cells as opposed to dead cells.

**Figure 3.**
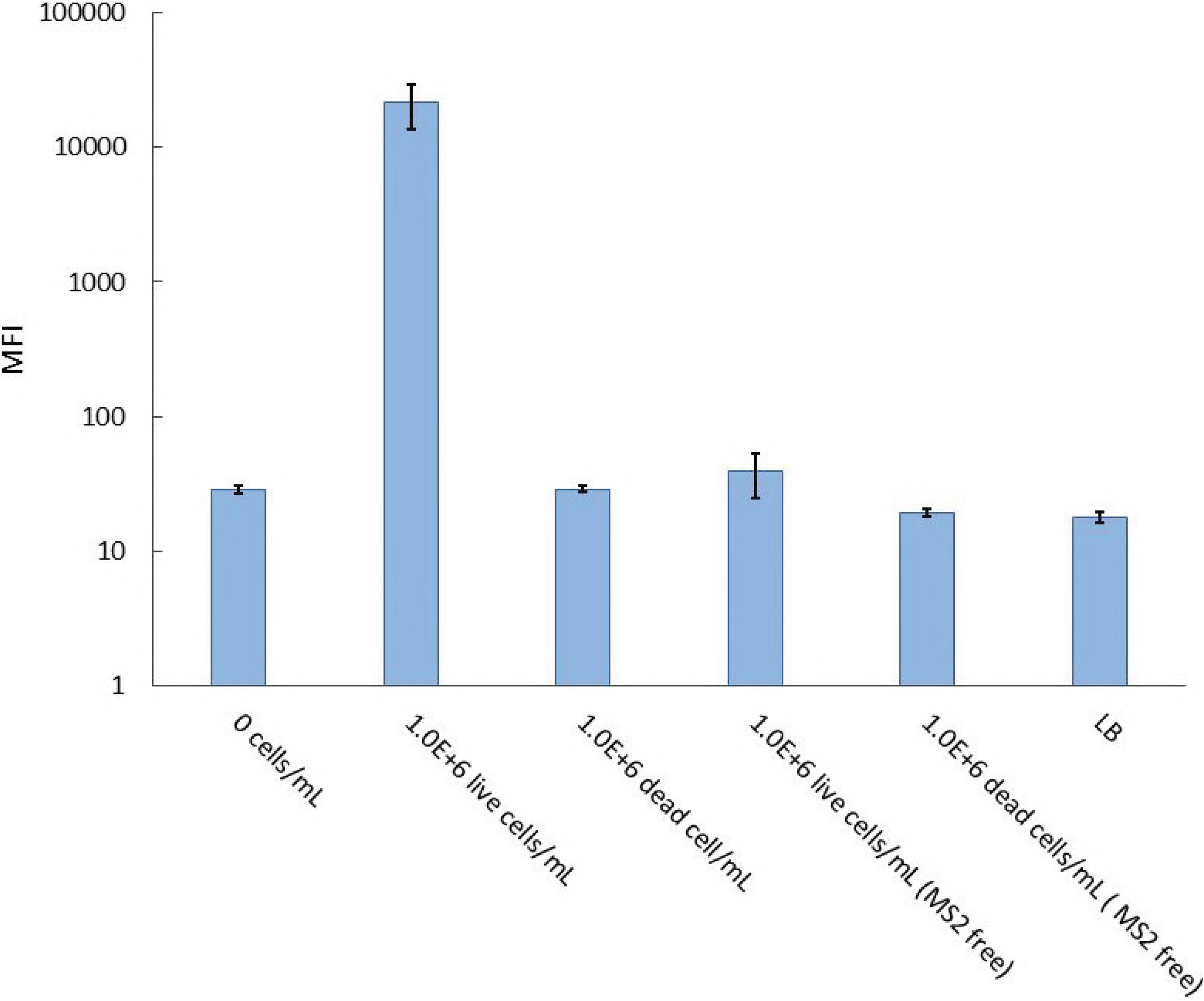
Validation of live cell discrimination ability of MS2 amplification coupled assay MS2 amplification-coupled MAGPIX immunoassay for *E. coli* detection was performed. Samples were incubated for 18 hours prior to analysis on the MAGPIX instrument. Vertical axis shows the median fluorescence intensity (MFI). The data in this figure is based upon 2 separate run of 2 replicates.

### Establishing the limit of detection and incubation time for the MS2 MAGPIX assay

Having established the live bacterial detection using the MS2 MAGPIX assay, next we investigated the sensitivity (limits of detection of *E. coli*) of the assay, and the minimal incubation time required for making a positive call in the assay. Signal amplification (as indicated by MFI) of the assay in the presence of varying concentrations of live *E. coli* (0-10^6^ cells/ml) was followed from 0 to 3 hours of incubation. The results indicated that with increasing concentrations and incubation times there was a corresponding increase in the signal intensities (Figure 4). Furthermore, the MFI was increased significantly in samples containing *E. coli* at 1×10^2^ cells/mL after 3-hour incubation and thus establishing a limit of detection for this assay. The limit of detection is defined by a signal greater than the mean background MFI plus three standard deviations. Higher concentrations of *E. coli* (10^5^-10^6^ CFU/ml) produced signal intensities that allowed detection in shorter incubation times; i.e., 1 hour where almost 9-fold increase was observed with 10^5^ cells/ml compared to 10^3^ cells/ml.

**Figure 4.**
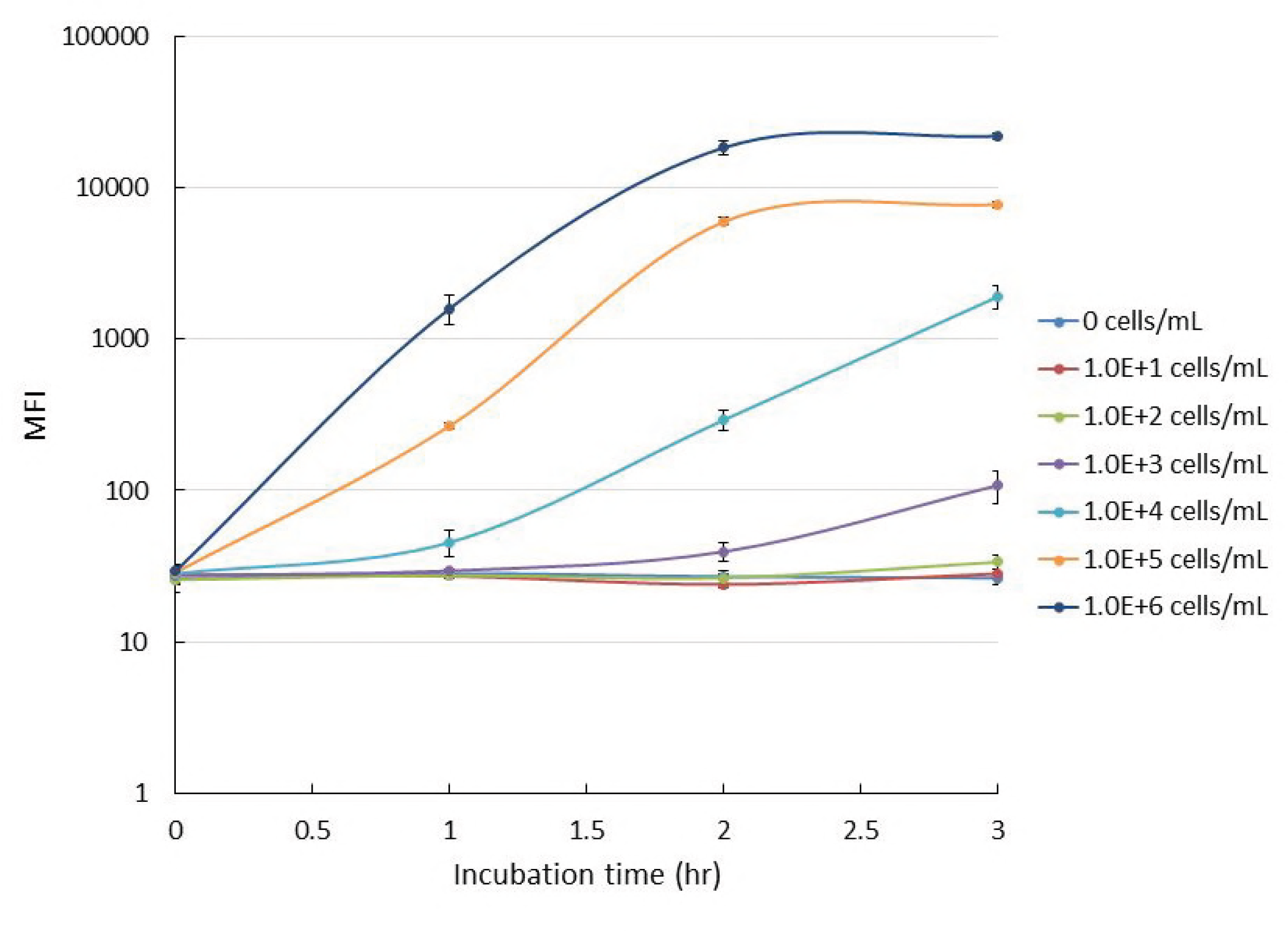
Determination of the limit of detection and incubation time for MS2 phage MAGPIX assay. Signal intensity of MS2 amplification-coupled MAGPIX immunoassay for *E. coli* detection was tracked from 0 to 3 hours. Samples containing 1.0 ×10^6^ PFU/mL of MS2 and live cells at varying concentrations from 0-1 x 10^6^ CFU/ml were incubated for indicated times prior to analysis on the MAGPIX instrument. Vertical axis shows the median fluorescence intensity (MFI) and each data point is the average of 2 replicates of 2 separate assays.

### Determination of MS2 phage binding selectivity between live and dead cells using MAGPIX assay

When bacterial samples are inactivated, it is difficult to assess complete inactivation by NASBA or immunoassay because dead bacteria still contain nucleic acids or immunoassay targets (epitopes) that are reactive to the respective assays. On the other hand, since phage amplification-coupled assay require live bacteria, it can be a valuable tool for validation of complete inactivation. However, incomplete inactivation may result in samples containing both live and dead cells. In this case, phage can potentially bind to both dead and live cells. Phage binding to dead cells can competitively inhibit binding to live cells (i.e., reducing the number of phages available for infecting live cells) and thereby reduce the sensitivity of the assay. To investigate if MS2 can selectively bind to live cells in the presence of dead cells, MS2 phage MAGPIX assay was performed in the presence varying concentrations of live cells (0-10^6^ cells/ml) or live and dead cells at the same concentrations (0-10^6^ cells/ml) for one hour. The results indicated that the signal intensities (as MFI) between the groups are very comparable indicating that the presence of dead cells did not affect the infection, replication and growth of live bacteria present in the sample (Figure 5). This result indicates that MS2 selectively bound to only living cells in the presence dead cells.

**Figure 5.**
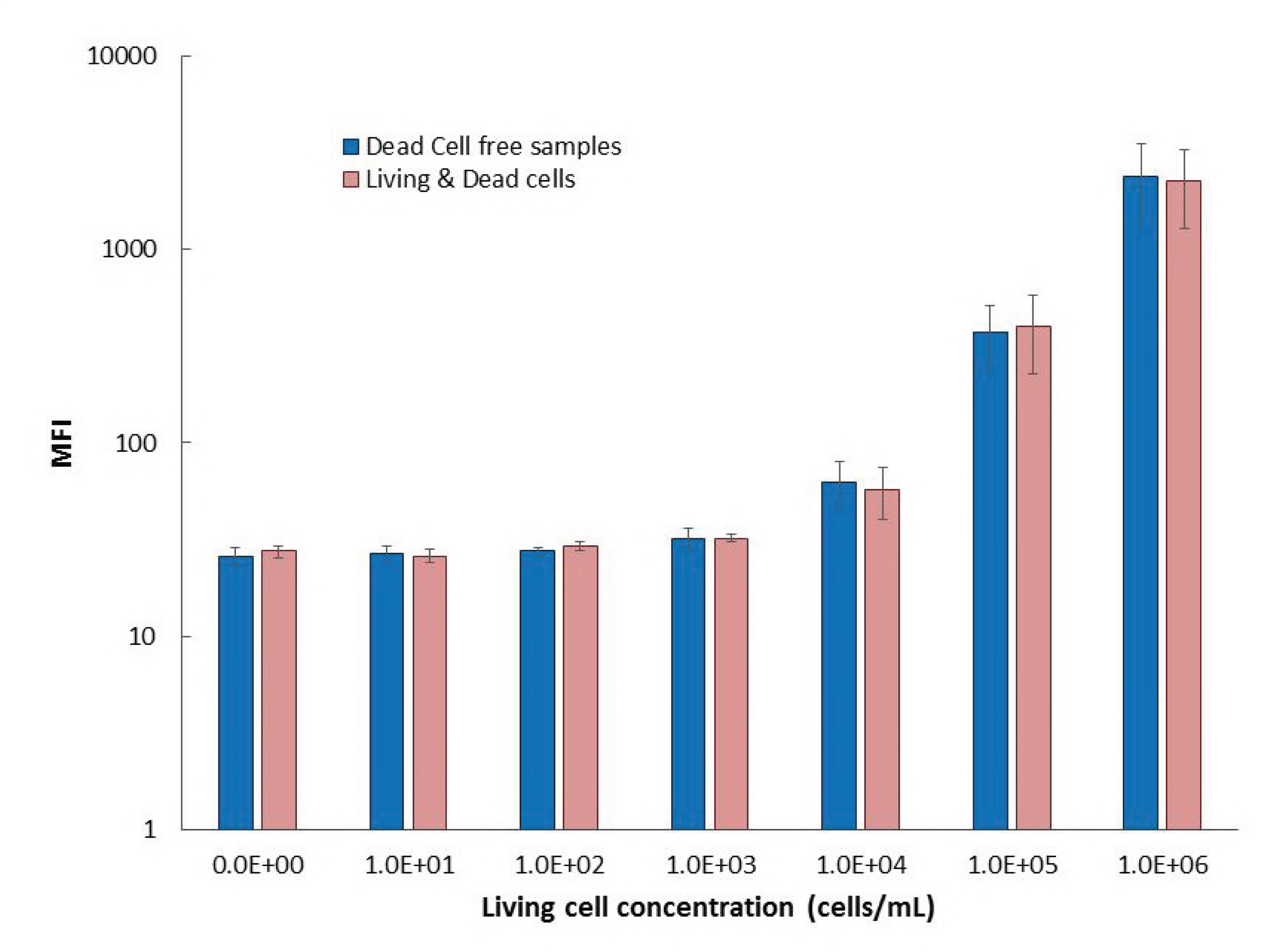
Determination of MS2 phage binding selectivity between live and dead cells using MAGPIX assay. Signal intensity of MS2 amplification-coupled MAGPIX immunoassay for *E. coli* detection was compared between sample containing only living cells and sample containing live cells and dead cells. Samples containing 1.0 ×10^6^ pfu/mL MS2 and live cells at respective concentration were incubated with/without 1.0 ×10^6^ cells/mL inactivated cells for 1 hour prior to the immunoassay. Vertical axis shows the median fluorescence intensity (MFI). The values represent average of two separate runs with 2 replicates each.

### Discussion

An ideal bioagent detection technology/platform would have the following desirable properties: The detection assays must have the potential for rapid, reproducible, high-sensitive (detection at very low concentrations of the agent), high-specific (high true positive/true negative and low false-positive/ low false-negative) detection of agents (conventional as well as uncharacterized or genetically modified agents) directly from complex matrix samples with minimal false results, capable of detecting low concentrations of target agents without interference from background materials. Also, the platform to conduct the assays should be user-friendly, portable and a point of care or field device that is capable of detecting multiple agents simultaneously in a high throughput manner (processing hundreds of samples) in any matrix type (clinical or environmental). Additionally it is highly desirable to have a flexible technology with an open architecture; amenable for a plug and play format to develop new assays rapidly and, above all, the technology should be inexpensive. Although several of the currently available commercial detection platforms provide many of the desired features, no one system can satisfy all of these criteria.

Many of the current rapid detection technologies are based either on PCR or some form of an immunoassay such as lateral flow immunoassay. The major drawback of these methods is that they cannot discriminate whether the suspected sample contains live or dead bacteria. This discrimination is critical especially in a biothreat/ biosurveillance scenarios not only for making correct courses of action but also for verifying if decontamination activities were successful. Also, in a clinical or point of care/field setting such a discrimination ability and phage mediated signal amplification will be very useful for determining antibiotic sensitivity rapidly. For example, in slow growing bacteria like *Mycobacterium* spp such an approach has been used to test antibacterial susceptibility in the field (11). Also, these technologies are limited in their potential for multiplexing and high throughput analysis. To address these two gaps, as a proof of concept, we developed a phage based live agent detection assay using MAGPIX instrument. Phage MS2 is an *E. coli* male specific phage because of its specificity to infect only strains that carry the F pilus (21). MS2 has a burst size of 5000-10000 per infected cell (22) and a short burst time of 30 minutes (23). In this study, MS2 amplification-coupled MAGPIX immunoassay was performed to demonstrate utility of bacteriophage amplification-coupled immunoassay for *E. coli* detection. Essentially, in this assay, progeny particles resulting from infection of bacteria in a sample are detected and reported as an indicator of the pathogen present. We found a sensitivity of 1 X10^4^ cells/ml of *E. coli* and selective binding to live cells and no inhibition in the presence of dead bacteria after a one hour incubation. This limit of detection is quite comparable to published immunoassays for *E. coli* detection (24, 25). The limit of detection of this assay is dependent on the burst size of a given phage in its specific bacterial host. Phages with high burst will yield large number of progeny particles that will yield correspondingly high signal intensity in the MAGPIX assay. However, for phages with low burst size, a large number of bacteria should be present during initial infection, in order to yield high enough signal intensity above the background to make a positive call. The optimal multiplicity of infection needs to be determined for each phage-bacterium combination since the initial bacterial load in each sample may vary. Intuitively, it would seem phage-based assays are not available for other types of agents such as spores, viruses and toxins. However, using M13 like phages, one can pan for phages with peptide displays that bind specifically to these biothreats and add that to the MAGPIX panel of assays. Despite these limitations, our future efforts will focus on developing a phage multiplex MAGPIX assay panel to target bacterial, viral and toxin threats.

## 2 Materials and Methods

### 2.1 Phage stock

MS2 phage and *E. col* strain C-3000 (ATCC 15597) used in this study were obtainedfrom Brouns lab, Delft University of Technology (NL). The phage stock was prepared from the seed stock in small scale liquid culture using established laboratory procedures (26)

### 2.2 MAGPIX immunoassay development

Polyclonal antibodies provided by Defense Biological Product Assurance Office (DBPAO); Rabbit anti-MS2 (Rab α-MS2: ABE#120, J-291100-02) were used to develop the capture sandwich assay on the MAGPIX platform (Luminex, Austin, TX). Antibodies were immobilized on MagPlex carboxylated microspheres (Luminex, Austin, TX) at 5pg antibody/ microsphere using the carbodiimide coupling protocol provided by the manufacturer. To conduct the coupling, 0.1 M Sodium Phosphate monobasic (pH6.2±0.2) was used for washing beads and incubation. Microspheres were then washed and resuspended in 0.01M PBS (pH7.4±0.1) with 0.05%Tween20. Antibodies immobilize on microsphere were used as the capture in the assay. Antibodies were also biotinylated using a 30-fold molar excess of BT-LCLC-NHS (Thermo Scientific, Rockford, IL) as the detection element with a streptavidin-phycoerythrin conjugate (SAPE) serving as the tracer molecule.

### 2.3 MAGPIX immunoassay procedure

2500 microspheres per well of antibody coated microspheres were incubated for 1 hour with samples. Following incubation, the sample was, washed, and then incubated with 4μg/mL biotinylated antibody for 30 minutes. The microspheres were washed again and then incubated with 4μg/mL SAPE to generate the fluorescent complex. After a final wash and re-suspending in 100μL of wash buffer, the assay results were evaluated using the MAGPIX instrument. Incubations were performed at room temperature, at 800rpm, and protected from light. Samples were washed twice with 100μL of wash buffer (0.01M PBS with 1%BSA and 0.05%sodium azide, pH7.4).

### 2.4 MS2 amplification-coupled MAGPIX immunoassays

MS2 amplification-coupled MAGPIX immunoassays were performed on the MAGPIX platform. Prior to MAGPIX immunoassay, MS2 and *E.coli* (Delft University of Technology, NL) diluted in LB media were incubated at 37 °C shaking at 300 rpm for respective time. E-coli concentration (cells/mL) was calculated by OD_600_ based on the formula as OD_600_ of 1.0 = 8 × 10^8^ cells/mL. To prepare near 100% of live cells, live E-coli used in this study was incubated to keep OD_600_ below 1.0 after inoculating. When dead cells were used, *E.coli* was inactivated by heating over 95°C in water bath for 15 min prior to the OD_600_ measurement. Incubated samples were analyzed by MAGPIX assay.

## Acknowledgements

We would like to thank Dr. Stan JJ Brouns and Franklin Nobrega (Delft University of Technology, Netherlands) for the kind gift of MS2 phage and the *E. coli* strain used in the assay. T. Mido was supported by the Japan Ministry of Defense – US Army Engineer and Scientist Exchange Program. Funding for this work was provided the Defense Biological Product Assurance Office, Joint Program Manager Guardian, Joint Program Executive Office for Chemical Biological Defense.

